# Improved assemblies and comparison of two ancient *Yersinia pestis* genomes

**DOI:** 10.1101/073445

**Authors:** Nina Luhmann, Daniel Doerr, Cedric Chauve

## Abstract

*Yersinia pestis* is the causative agent of the bubonic plague, a disease responsible for several dramatic historical pandemics. Progress in ancient DNA (aDNA) sequencing rendered possible the sequencing of whole genomes of important human pathogens, including the ancient *Yersinia pestis* strains responsible for outbreaks of the bubonic plague in London in the 14th century and in Marseille in the 18th century among others. However, aDNA sequencing data are still characterized by short reads and non-uniform coverage, so assembling ancient pathogen genomes remains challenging and prevents in many cases a detailed study of genome rearrangements. It has recently been shown that comparative scaffolding approaches can improve the assembly of ancient Yersinia pestis genomes at a chromosome level. In the present work, we address the last step of genome assembly, the gap-filling stage. We describe an optimization-based method AGapEs (Ancestral Gap Estimation) to fill in inter-contig gaps using a combination of a template obtained from related extant genomes and aDNA reads. We show how this approach can be used to refine comparative scaffolding by selecting contig adjacencies supported by a mix of unassembled aDNA reads and comparative signal. We apply our method to two data sets from the London and Marseilles outbreaks of the bubonic plague. We obtain highly improved genome assemblies for both the London strain and Marseille strain genomes, comprised of respectively five and six scaffolds, with 95% of the assemblies supported by ancient reads. We analyze the genome evolution between both ancient genomes in terms of genome rearrangements, and observe a high level of synteny conservation between these two strains.

## 1 Introduction

*Yersinia pestis* is the pathogen responsible for the bubonic plague, a disease that marked human history through several dramatic pandemics, including the Justinian Plague and the Black Death. It diverged a few thousands years ago from a relatively non-virulent pathogen, *Yersinia pseudotuberculosis*. The precise timing of the divergence between these two pathogens is still controversial^40^, but it is widely accepted that the emergence of *Yersinia pestis* as a virulent human pathogen was characterized, among other elements, by the acquisition of numerous repeat sequences, especially Insertion Sequences (IS) that triggered an extensive chromosomal rearrangement activity^9, 11^. Also worth noting, loss-of-function mutations that can be due to chromosomal rearrangements have been identified as evolutionary adaptation for flea-borne transmission from *Yersinia pseudotuberculosis* in the ecological context^21^. This makes the *Yersinia* family appear as an interesting model for the study of genomes rearrangements during pathogen evolution.

Traditionally, the study of genome rearrangements relies on a comparative approach using the genomes of related extant organisms. Under appropriate models of evolution, this comparison provides indirect insight into genomic features of ancient species and their evolution toward extant species, see^11^ for example for the specific case of genome rearrangements in *Yersinia*. However, this approach requires well assembled extant genomes, as otherwise it is difficult to distinguish breakpoints due to assembly fragmentation from evolutionary breakpoints. For example, Auerbach *et al*^*2*^ discuss several chromosomal rearrangements between two closely related *Yersinia pestis* strains, but could not determine the evolutionary history of these modifications as related strains are only partially assembled and highly rearranged. Besides challenges for the analysis of genome rearrangements, fragmented assemblies of bacterial genomes impede subsequent analysis like genome annotation, the identification of gene duplication, gene loss and lateral gene transfer, or the characterization of gene families, as well as the analysis of intergenic and especially repeat-rich genomic regions which are usually not assembled^17, 24, 35, 48^. Finally, while synteny breakpoints often coincide with gaps in a conservative assembly, unfinished assemblies also pose the jeopardy of uncorrected mis-assemblies influencing the reconstruction of genome rearrangement events^37, 47^.

In contrast to the approach based on comparing extant genomes, sequenced ancient DNA (aDNA) extracted from conserved remains can give direct access to the sequence of ancient genomes and thus, theoretically, allows us to study the evolution from ancestors to descendants directly. Following advances in aDNA high-throughput sequencing technologies and protocols^18, 20, 22, 33, 34, 52^, the genomes of several ancient human, animal and plant pathogens have recently been sequenced at the level of complete or almost complete chromosomes, including the agents of potato blight^31, 53^, brucellosis^23^, tuberculosis^5^, leprosis^44^, *Helicobacter pylori*^30^, cholera^12^ and of the bubonic plague^6, 7, 50^, leading to important historical and evolutionary discoveries. However, unlike extant DNA high-throughput sequencing that is experiencing a breakthrough transition towards long-reads, aDNA sequencing methods generate extremely short reads with low and non-uniform coverage^52^. As a result, aside of rare exceptions^44^, the assembly of aDNA reads generates numerous short contigs. For example, a reference-based assembly of the Black Death pandemic agent resulted in several thousand contigs^7^, two thousand of them of length 500bp and above. While short aDNA reads can be mapped onto one or several extant reference genomes to detect important evolutionary signals such as SNPs and small indels^36, 43^, they lead to fragmented assemblies which makes it challenging to exploit aDNA sequencing data similar to fragmented assemblies of extant strains to analyze the evolution of pathogen genome organization.

Without long-read sequencing data, comparative scaffolding based on the comparison of the contigs of a genome of interest with related assembled genomes has proven to be a useful approach to improve the assembly of fragmented genomes, especially bacterial genomes^8, 25, 39, 41^. Among such methods, FPSAC^39^ was introduced to improve ancient genome assemblies within a phylogenetic context. It was applied to aDNA contigs from the *Yersinia pestis* strain responsible for the medieval London bubonic plague outbreak – that was shown to be ancestral to several extant *Yersinia pestis* strains^7^ – and resulted in an improvement of the initial contig assembly from thousands of contigs to a chromosome-scale scaffolding. Moreover, taking advantage of the high sequence conservation in *Yersinia pestis* genomes, the inter-contigs gaps of the ancient *Yersinia pestis* strain were filled with putative sequences reconstructed from multiple sequence alignments of conserved extant gaps. This gap-filling step shed an interesting light on genomic features hidden within the assembly gaps, in particular IS and their correlation with rearrangement breakpoints reuse, but also allowed the potential reconstruction of regions that were not recovered or were absent from the aDNA material. However, the scaffolding of adjacencies and gap sequences obtained in^39^, that accounted for roughly 20% of the genome size, were inferred through computational methods within a parsimony framework, that can be sensitive to convergent evolution that cannot be ruled out for genomes with a high rate of genome rearrangements such as *Yersinia pestis*^11^.

In the present work, we address this issue by using the large set of aDNA reads that are unassembled after the contig assembly stage, to confirm the scaffolding of contigs as well as sequences for inter-contigs gaps. We introduce the method AGapEs (Ancestral Gap Estimation) which attempts to fill the inter-contig gap between two adjacent ancient contigs by selecting a set of overlapping aDNA reads that minimizes the edit distance to a template gap sequence obtained from the extant genome sequences that support the adjacency. We directly include annotations of potential Insertion Sequences (IS) in the extant genomes in the analysis to use the aDNA reads when the presence of an IS in the ancient genome is doubtful due to a mixed signal of presence/absence in the supporting extant genomes.

We apply this strategy to two data sets of ancient DNA reads for ancestors of the human pathogen *Yersinia pestis*^9, 32^. This bacterium is the causative agent of the bubonic plague and responsible for three major epidemics, the last one still on-going. The first aDNA data was obtained from London victims of the Black Death pandemic in the 14th century^7^, and the second consists of five samples from victims of Great Plague of Marseille around 400 years later^6^. For both data sets, we obtain an assembly with reduced fragmentation and are able to fill a large number of inter-contig gaps with aDNA reads. We identify several genome rearrangements between the ancient strains and extant *Yersinia pestis* genomes, however observe only a single small inversion between both ancient strains, suggesting that the genome organization of the agent of the second major plague pandemic was highly conserved.

## 2 Materials and Methods

We first describe the input to our analysis, namely ancient sequencing data, ancient and extant assemblies and annotations of IS, before outlining the general pipeline we used to improve the assembly of the ancient genomes.

### Sequencing data and reference genomes

The first aDNA data set was obtained from a London victim of the Black Death pandemic in the 14th century^7^ (individual 8291), the second consists of five samples from victims of Great Plague of Marseille around 400 years later^6^. The average read length is 53 bp in the London dataset and 75bp in the five Marseille samples (Figure S3). We rely on seven extant *Yersinia pestis* and four *Yersinia pseudotuberculosis* as reference and outgroup genomes (see Table S1). The phylogeny of the considered strains is depicted in Figure S1 and is taken from^6, 7^.

### Contig assembly and preprocessing

We de novo assembled aDNA reads into contigs using Minia^10^ for both aDNA data sets (London outbreak and Marseille outbreak). Minia is a conservative assembler based on an efficient implementation of the de Bruijn graph methodology. In general, Minia produces shorter contigs than competing assemblers, as it avoids assembly decisions in case of ambiguity in the sequence data. We will refer to the Minia assemblies as *de novo* assemblies in the following. To allow the comparison with extant genomes, contigs above a minimum length threshold were aligned with the extant genomes to define families of homologous synteny blocks (called markers from now) as described in^39^. Marker families were then filtered to retain only one-to-one orthologous families, i.e. families that contain one and exactly one marker in each considered extant and ancient genome.

### Insertion sequence annotation

Insertion Sequences (IS) are strongly related to rearrangements in *Yersinia pestis* evolution, and their annotation in the considered extant genomes is crucial. In order to annotate IS, we designed our own annotation pipeline. Because IS elements in the original Genbank files were rather disparately annotated, we relied on automated annotations from the Basys annotation server^49^. Basys identified 11 families of IS transposase proteins (see Table S2). For each of these families, we produced a multiple alignment of their annotated sequences using `muscle`^15^ which was subsequently used to train Hidden Markov Model (HMM) profiles. Using `hmmer`^14^, we then annotated those regions as associated to IS elements that showed significant correlation to any of the HMM profiles. We eventually combined the Genbank annotations with these derived annotations. The number of these IS annotations per reference genome ranges from 151 in *Yersinia pestis* KIM10+ to 293 in *Yersinia pestis Antiqua* (see Table S1). The length of the annotations ranges from 60bp to 2,417bp; some short annotations deviate from the expected length for IS, however, in order to avoid filtering any true annotations, we include them all as potential IS coordinates in the downstream analysis.

### Ancestral marker adjacencies

Each marker can be defined by a pair of marker extremities. An adjacency consists of two markers extremities that are contiguous along a genome, *i.e.* are not separated by a sequence containing another marker. For extant genomes, extant adjacencies can be observed directly, while for an ancestral genome of interest, we infer potential ancestral adjacencies using the Dollo parsimony principle^39^: two ancient marker extremities are potentially adjacent if there exist two extant genomes whose evolutionary path contains the most common recent ancestor of the London and Marseille strains and where the two corresponding extant marker extremities are contiguous (see Figure S5 for an example). Hence every potential ancestral adjacency is supported by a set of extant adjacencies. A gap is the sequence between the two marker extremities defining an adjacency. Therefore each putative ancestral gap is likewise supported by a set of extant gap sequences.

We say that two potential ancestral adjacencies are *conflicting* if they share a common marker extremity. An *IS-annotated* adjacency is supported by at least one extant adjacency whose gap contains an IS annotation. An adjacency that is neither conflicting nor IS-annotated is said to be *simple*.

### AGapEs: Assembly of ancestral gap sequences from aDNA reads

The main methodological contribution we introduce is a template-based method to assess the validity of a potential ancestral adjacency. The general principle is to associate to every ancestral gap a template sequence obtained from the supporting extant gaps sequences. We can then map aDNA reads onto this template and assemble the mapped aDNA reads into a sequence that minimizes the edit distance to the template sequence. The rationale for this template-based approach is that, due to the low coverage of the aDNA reads and their short length, existing gap-filling methods fail to fill a large number of ancestral gaps. For example, the method gap2Seq^42^, a recent efficient gap-closing algorithm based on finding a path of given length in a de Bruijn graph, is not able to fill roughly half of the ancestral gaps of the Black Death data set (see Table S7).

We describe now the AGapEs algorithm. Assume we are given a template sequence t for a gap in an adjacency a = {m_1_, m_2_} between two marker extremities. We define *R* = *m*_1_ +*t* + *m*_2_ as the concatenated nucleotide sequence of the oriented markers and the respective template. We first align the aDNA reads onto R, using BWA^29^, where we only consider mappings whose start and/or end position is in t (i. e. either fully included in t or overlapping the junction between a marker and the gap template). Next, we construct a graph *G* where vertices are mappings 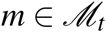 and there is an edge between two vertices (i. e. mappings) if the two mapping coordinates (segments of *R*) overlap. For each such edge/overlap, we define s as the non-overlapping suffix of the mappings with the highest end coordinate. We can then associate a weight to each edge given by the edit distance between s and the subsequence *R_s_* of *R* it aligns to. A sequence of overlapping reads that minimizes the distance to *t* can then be found by searching for a shortest path between the vertex labeled with the smallest start position (i. e. the first mapping covering the junction between *m*_1_ and *t*) and the vertex labeled with the largest start position (i. e. the last mapping covering the junction between *t* and *m*_2_). See Figure S8 for an illustration.

If such a path exists, it can be found with Dijkstra’s algorithm^13^ implemented based on a min-priority queue in O (|*E*| + |*V* | log |*V* |) time, where *V* is the vertex set and *E* the edge-set of *G*. If no such path exists, then there are either regions in *R* that are not covered by any mapped aDNA read or breakpoints in the mapping, where two consecutive bases in the sequence are covered, but not both by the same read. In these cases, uncovered regions and breakpoints need to be identified in the mapping beforehand to identify start and end vertex of the shortest path. We can then obtain a partial gap filling, precisely for the regions covered by mapped reads.

The key element of the approach described above lies in defining the template sequence or set of alternative template sequences associated to each ancestral gap. We follow the general approach described in^39^, that computes a multiple sequence alignment of the supporting extant sequence gaps and applies the Fitch-Hartigan parsimony algorithm^16^ to each alignment column to reconstruct a most parsimonious ancestral sequence. If the multiple sequence alignment of extant gaps shows little variation, as is the case for most gaps in our data sets, then a single template sequence can be considered, as we expect that minor variations compared to the true ancestral sequence (substitutions, small indels) will be corrected during the local assembly process outlined above. Alternatively, if larger variations are observed, such as larger indels or a contradicting pattern of presence/absence of an IS in the supporting extant gaps, then alternative templates can be considered, under the hypothesis that the true variant can be recovered from the mapped aDNA reads.

Hence in the following analysis, we separate all potential ancestral gaps into groups of simple, conflicting and IS-annotated gaps. For simple and conflicting gaps without IS annotation, we can follow the process described above directly. For IS-annotated gaps, we reduce the described large variations in the multiple alignment by further dividing its supporting extant gaps into sets of IS-annotated and non-IS-annotated sequences respectively. Building the multiple alignment on each of these sets separately allows us to define two alternative templates that can be used as a basis to fill the gap. Ideally, differences in read coverage or breakpoints naturally identified by AGapEs then point to one of the alternative templates for each IS-annotated gap. Further, for each template that is only partly covered by mapped reads, we will correct the covered parts according to the reads using AGapEs and use the template sequence otherwise.

The implementation of AGapEs is available at http://github.com/nluhmann/AGapEs, the data underlying the following results can be downloaded from http://paleogenomics.irmacs.sfu.ca/DOWNLOADS/AGAPES_data_results.zip.

## 3 Results and Discussion

### 3.1 The London strain

The de novo assembly consists of 4,183 contigs of length at least 300bp that cover 2,631,422 bp (see Suppl. Material A.4). Using the marker segmentation described in the previous section, we subsequently obtain 3,691 markers covering 2,215,596 bp in total. Note that not all contigs are represented in the marker set, as no part of these contigs aligns uniquely and universally to all reference genomes.

#### Reconstructing potential ancestral adjacencies

We obtain 3,691 potential ancestral adjacencies: 3,483 are simple, 201 are IS-annotated and non-conflicting, and only 7 are conflicting. Among the conflicting adjacencies 5 are also IS-annotated, illustrating that most rearrangements in *Yersinia pestis* that can create ambiguous signal for comparative scaffolding are associated with IS elements (see also Table S5).

For most potential ancestral adjacencies, the lengths of the sequences in extant genomes associated with the supporting extant adjacencies are very similar, indicating well conserved extant gaps (Figures S7(a) and S7(b)). There are 21 gaps whose lengths difference falls into the length range of potential annotated IS elements, thus raising the question of the presence of an IS within these adjacencies in the ancestral genome. We note a small number of 5 potential ancestral adjacencies with strikingly large extant gap length differences. All of these gaps accumulate more than one IS annotation in some extant genomes. Most problematic is a gap with length difference of more than 100.000 bp. As this gap is not well conserved in general (apart from the inserted sequences), it is difficult to obtain a good template sequence based on a very fragmented multiple alignment. We will get back later to this special gap.

#### Ancestral gaps filling

We applied AGapEs to all potential ancestral gaps. We assume a gap to be filled, if we find a sequence of reads that covers the whole ancestral gap. As we test two alternative templates for an IS-annotated gap, we consider it filled if only one alternative is covered or if both templates are covered but the IS is only annotated in a single extant genome. In the latter case, we expect the non-IS gap version to be ancestral, as the IS most appeared along the edge to the annotated extant genome. If otherwise both alternative template sequences are covered, we cannot recover the true positive gap at this point and mark it as not filled. If a gap template sequence is only partially covered by mapped aDNA reads, we correct the covered regions as described above and use the template sequence of the uncovered regions to complete filling the gap. Figure 1 summarizes the gap-filling results (see also Table S5).

**Figure 1.**
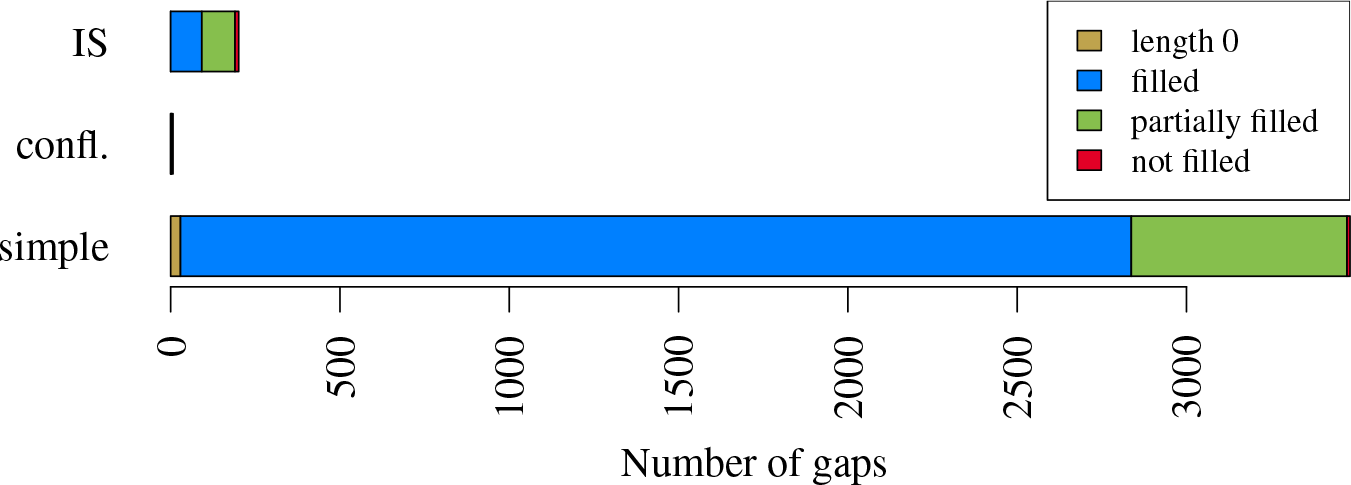
Result of gap filling for the London data set. Note that if a gap is conflicting and IS-annotated, we assign itto the conflicting group. We differentiate between gaps of length 0 (i. e. both markers are directly adjacent), completely and partially filled gaps, and not filled gaps.

A high number of gaps is supported by sufficient read coverage that enables us to fill the gap with a sequence of overlapping aDNA reads. Especially considering partially covered gaps improves the length of the genome that is supported by reads. Note that we also find covering reads for all gaps of length 0, spanning the breakpoint between directly adjacent markers.

We further computed the edit distance between each reconstructed gap sequence and its previous gap template. For IS-annotated gaps, we computed the distance to a template sequence based on all extant gap occurrences, i. e. without considering alternative templates as described previously. We identified one case where the parsimonious gap sequence based on all extant occurrences of the adjacency excludes the IS. However if aDNA reads are mapped separately to alternative templates based on IS and non-IS annotated extant gaps, only the IS-annotated gap template is covered.

For IS-annotated gaps, 95 ancestral gaps contain an IS, while 106 ancestral gaps are reconstructed without the IS. From these 95 IS gaps, 22 contain annotations that are shorter than 400bp, however they all contain additional longer annotations in the same gap. Analyzing the number of ancestral IS with a Dollo parsimony criterion considering only the extant IS annotations, we have 96 ancestral gaps that contain an IS, indicating a large agreement between the IS that are conserved by the parsimony criterion and the IS supported by aDNA reads.

#### Conflicting adjacencies

Conflicting adjacencies are related by the marker extremities they share, defining clusters of related conflicting adjacencies. We identified two such clusters (see Figure S11). One consists of three adjacencies that are all annotated with IS elements, while the other consists of four adjacencies, including two IS-annotated adjacencies. In total, only two of these conflicting adjacencies are supported by aDNA reads. All other adjacencies contain uncovered regions indicating potential breakpoints. So in order to propose a conflict-free scaffolding, we chose to remove all unsupported conflicting adjacencies. See Figure S12 for the read coverage of discarded adjacencies. Note that filling these gaps only partially does not provide much information, as uncovered regions can be either breakpoints or correspond to regions of the ancestral genome that were not sequenced.

The set of ancestral adjacencies can then be ordered into five Contiguous Ancestral Regions (CARs). We converted the reconstructed sequences of markers back to genome sequences by filling the gaps with the read sequences if possible and resorting to the template sequence otherwise.

As mentioned earlier, we observe one gap with highly differing extant gap lengths and very little conservation in the reconstruction. The multiple alignment based on extant gap sequence is very fragmented and the mapping of reads onto this template is poor: the gap contains 211 uncovered regions of 9,319 bp in total. See Figure S13 for an overview over the read coverage for this gap in the de novo assembly. As the reconstructed sequence has a high edit distance after partial gap filling, we remove this gap sequence completely at this point to avoid dubious and non-robust reconstructed ancestral sequences.

In addition, we aligned all reads again to the final assembly to assess the amount of uncovered regions in the reconstructed sequences. In total, 88,529bp are not covered by any read, however most uncovered regions are rather short (see Figures S16 and S17). Based on this mapping, we ran the assembly polishing tool Pilon^51^ on the final assembly. It identified several positions where the assembled base (also present in the template) is the minority in comparison to all reads mapping at this position. As Pilon is not taking the respective bases of the extant genomes into account, it runs the risk of correcting the assembly according to sequencing errors in the reads. In fact, the most frequent proposed substitutions correspond to the common damage pattern of cytosine deamination observed in aDNA^34^. As a consequence, we only keep small indel corrections by Pilon but reject all single-base corrections.

In the improved assembly, 49.88% of the sequence is based on markers and hence directly adopted from the initial assembly. Together with the gaps that have been filled by read sequences, we can say that in total 95.25% are reconstructed using only the available aDNA reads.

#### 3.2 The Marseille strain

This data set consists of five samples as described in^6^ that we assembled separately with Minia^10^. We first compared the quality of the resulting assemblies by mapping contigs with a minimal length to the genome of the extant strain *Yersinia pestis CO92* and summing the total length of the mappings as seen in Figure 2. While restricting the minimal contig length, two of the samples cover an extensively larger part of the CO92 strain genome, and thus indicate a better sequencing quality. Figure 3 shows that if we restrict the minimal contig length, only a small part of the *Y*. *pestis* reference genomes is covered by contigs from all five samples.

**Figure 2.**
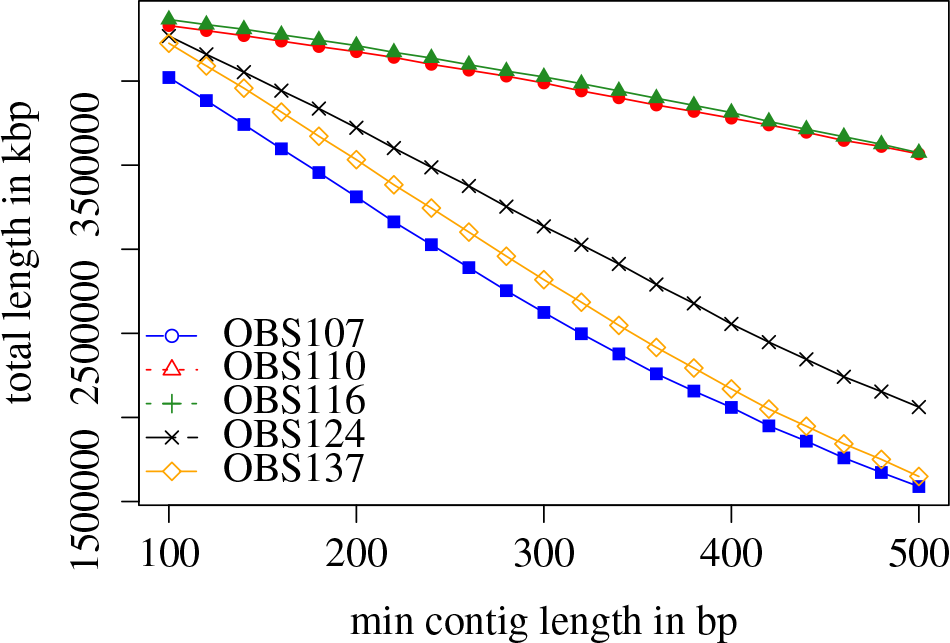
Total length of contigs mapped to*Yersinia pestisCO92* greater than a minimum contig length.

**Figure 3.**
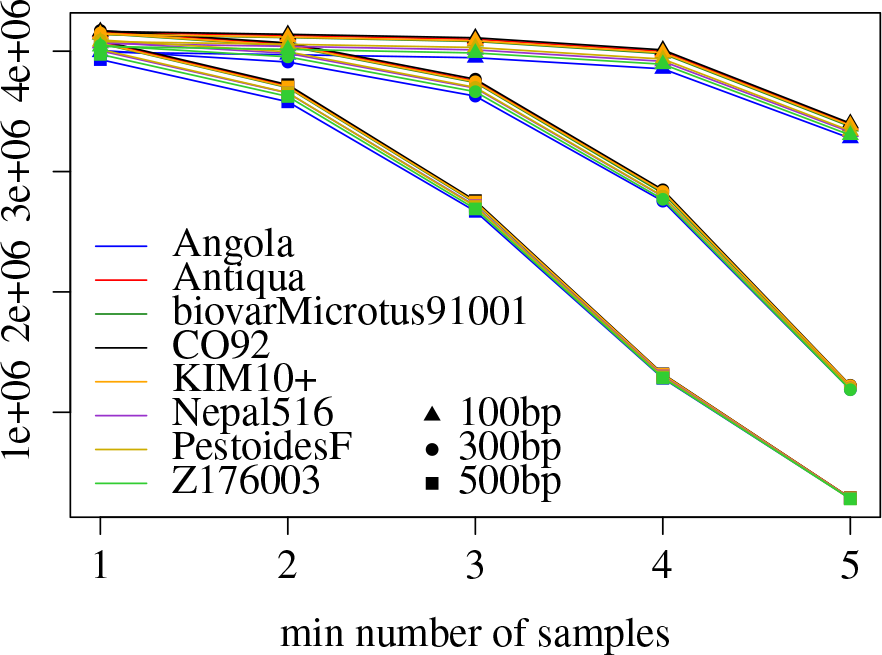
Comparison of assembled contigs by mapping to different reference sequences. While most of the references are covered by at least one sample, only a small part of the references is covered by all five samples.

We used the assembly of sample *OBS116* with a minimal contig length of 500bp to segment the extant genomes into markers. The assembly consists of 3,089 contigs with a total length of 3,636,663bp. The segmentation results in 2,859 markers with a total length of 3,143,627bp. We analyze 2,859 potential adjacencies: 27 of these gaps have a length of 0, leaving 2,832 gaps to fill. Based on the observations above, we joined all sample reads sets for filling the gaps in the reconstruction to achieve a better coverage.

We can see in Figure 4 that with the combined set of reads, we can fill nearly all simple gaps by read sequences. In addition, we obtain a higher number of IS-annotated gaps that are filled in comparison to the London data set. For the IS-annotated gaps, 95 are reconstructed containing the IS, 21 contain IS annotations shorter than 400bp. Hence we identified the same number of potential ancestral IS as for the London strain.

**Figure 4.**
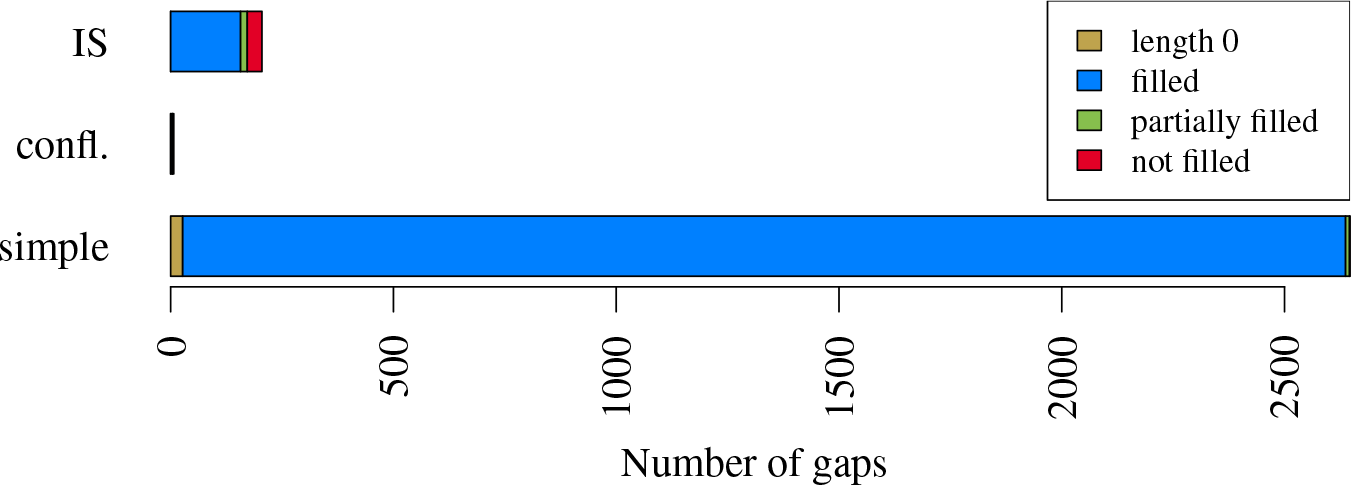
Result of gap filling for the Marseille dataset. Note that if a gap is conflicting and IS-annotated, we assign itto the conflicting group. We differentiate between gaps of length 0 (i. e. both markers are directly adjacent), completely and partially filled gaps, and not filled gaps.

We identified two conflicting components in this set of potential adjacencies (see Figure S14). Both of them align in terms of gap lengths and extant occurrences with the two components observed in the assembly for the London strain. In the first component, again only one conflicting adjacency is covered by reads. However, this is a different adjacency in comparison to both reconstructions for the London strain, while on the other hand we have no read support for the gap that is covered in the London data set. This could indicate a potential point of genome rearrangement (see discussion in next section). In the second component, all involved adjacencies are covered by reads from the five samples. In order to obtain a set of high confidence ancestral CARs, we removed all conflicting adjacencies in this component from the set of potential adjacencies. The coverage of all discarded adjacencies is shown in Figure S15.

This results into 6 CARs for the ancestral genome. Again, we used *BWA*^29^ to align reads from all five samples again to the assembly to assess the amount of uncovered regions in the reconstructed sequences. In total, only 54,672bp in this mapping are not covered by any read and the length of the uncovered regions is rather short (see Figure S17).

#### 3.3 Comparison of the London and Marseille strains genomes

As the Marseille *Yersinia pestis* strain is assumed to be a direct descendant of the London Black Death strain^6^, we aligned the obtained CARs in both reconstructions to identify genome rearrangements. As shown in Figure 5, apart from one larger deletion and one larger insertion in the Marseille strain related to the removed gap sequence in the London strain and a small inversion of length 4,138bp marked in black, the reconstructed CARs show no larger rearrangements between both genomes (grey links).

**Figure 5.**
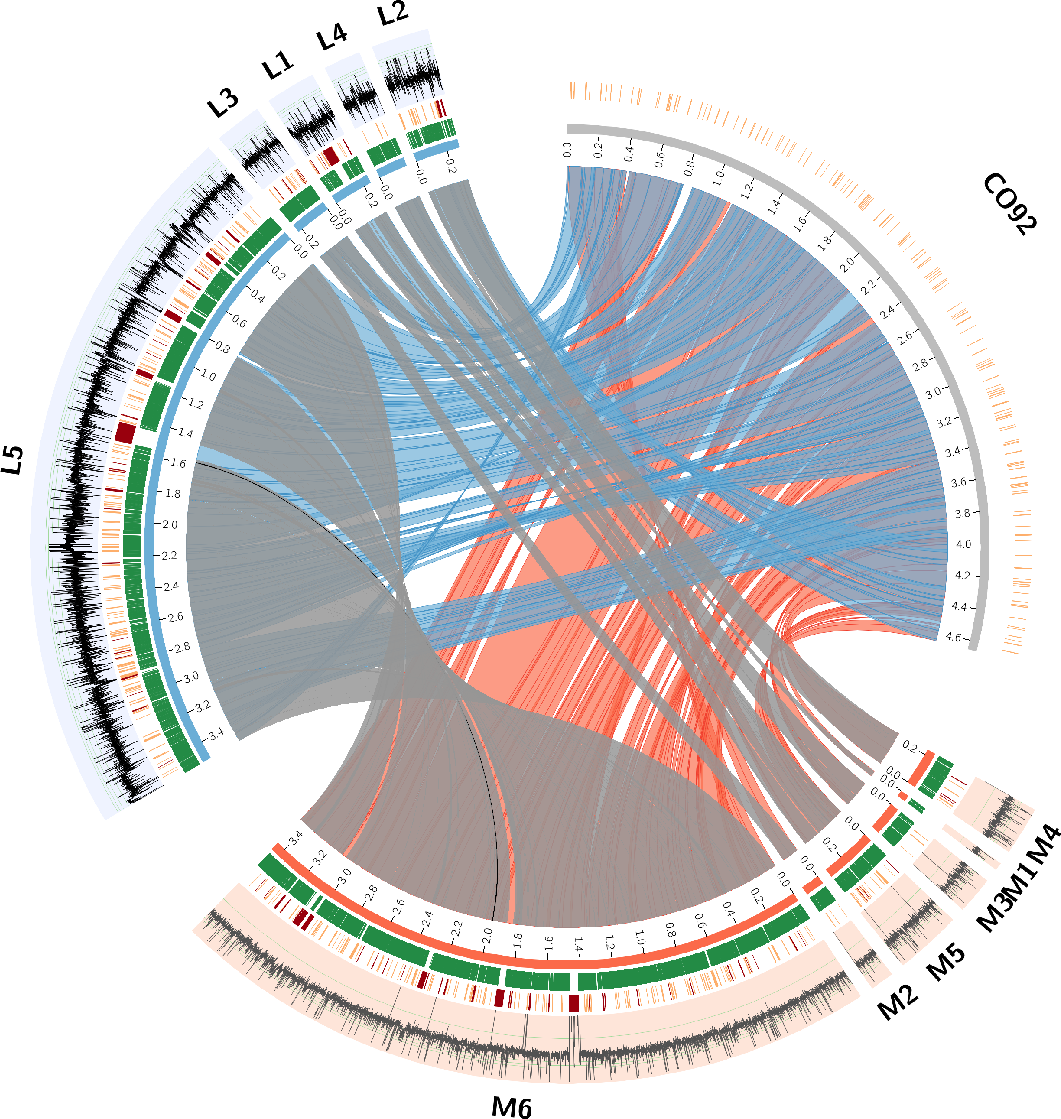
Comparison between the de novo assembly of the London strain (blue) and the Marseille strain (red) with the reference *Yersinia pestis CO92*. The inner links connect corresponding CARs in the reconstructions and the reference. Note that there is only a small inversion marked in black among the grey links. The positions in both reconstructions covered by markers are indicated in green. All gaps that have IS annotations in the extant genomes are shown in orange. For CO92, all IS annotations are shown as well. In addition, gaps that are only partially filled or have very unconserved extant gap lengths are indicated in red. Finally, the most outer ring shows the average read coverage in windows of length 200bp in log scale. Figure done with Circos^27^.

The difference in conflicting adjacencies kept is a possible indication for a rearrangement that however cannot be explicitly identified at this point. It causes the split pattern observed between CAR3 and CAR1 in the London strain and CAR2 and CAR5 in the Marseille strain. Given that the available read data does not allow us to further order the resulting CARs into a single scaffold, additional potential rearrangements could be assumed to be outside of the reconstructed CARs. In contrast, Figure 5 depicts several inversions and translocations between both ancient sets of CARs and the extant *Yersinia pestis CO92* (red and blue links respectively).

To clarify this further, we computed all potential orderings for both sets of CARs and determined the Double-Cut-and-Join (DCJ) ^4^ genome rearrangement distance (see Suppl. Material 1) for all such orderings between both ancient strain as well as in comparison to *Yersinia pestis CO92*. We obtain a weighted average distance of 4.04 between both ancient strains and an average distance of 11.16 to *CO92* with a standard deviation of 0.89 and 0.83 respectively. This suggests a much slower evolution in terms of rearrangements between both ancient strains and the extant strain.

### 3.4 Assembly evaluation

#### Influence of initial assembly

Bos et al^7^ describe a reference-based assembly of the London strain consisting of 2,134 contigs of length at least 500bp. It was obtained with the assembler Velvet^54^ using the extant strain *Yersinia pestis CO92* as a reference. In order to assess the influence of the reference sequence in the assembly of the ancient genome, we compare our pipeline using this initial assembly to our results based on the de novo assembly.

We compared the two sets of CARs obtained from both initial assemblies by aligning the resulting genome sequences using MUMmer^28^. We observe no rearrangements between both resulting sets of CARs (see also Figure S16), showing that, in terms of large-scale genome organization, the final result does not depend on the initial contig assembly.

#### Assembly validation

We compare our results to assemblies obtained with several other assembly pipelines. We used the *iMetAMOS* pipeline^26^ to determine the best de novo assembly for both data sets testing different assemblers. The winning assembly computed by SPAdes^3^ for both data sets as well as the minia assemblies on both datasets were subsequently used as input for two comparative scaffolding programs, Ragout^25^ and MeDuSa^8^, to obtain a scaffolding of the initial contigs considering the extant reference genomes. For all scaffolds, we ran gap2Seq^42^ to close the gaps. We will distinguish the results according to the scaffolding tool used in the following.

As shown in Table 1, for both datasets, Ragout is reconstructing the smallest number of contigs, however the scaffolds still contain a high number of unfilled gaps that cannot be closed by gap2Seq. See Table S10 for the results of all tool combinations. Our AGapEs reconstruction - although slightly more fragmented - achieves the best assembly likelihood according to both the LAP^19^ and CGAL^38^ score. The MeDuSa scaffolder is not able to estimate gap sizes as needed as input for gap2Seq, hence the better likelihood in comparison to Ragout can be accounted to the missing gaps characterized by N’s in the Ragout assembly. Also worth noting with the Marseille strain, MeDuSa was not able to correct a larger than expected contig assemblies obtained with SPAdes. Finally, the Minia-AGapEs assemblies do not contain Ns due to the filling of the gaps uncovered by reads by the template sequence.

**Table 1.**
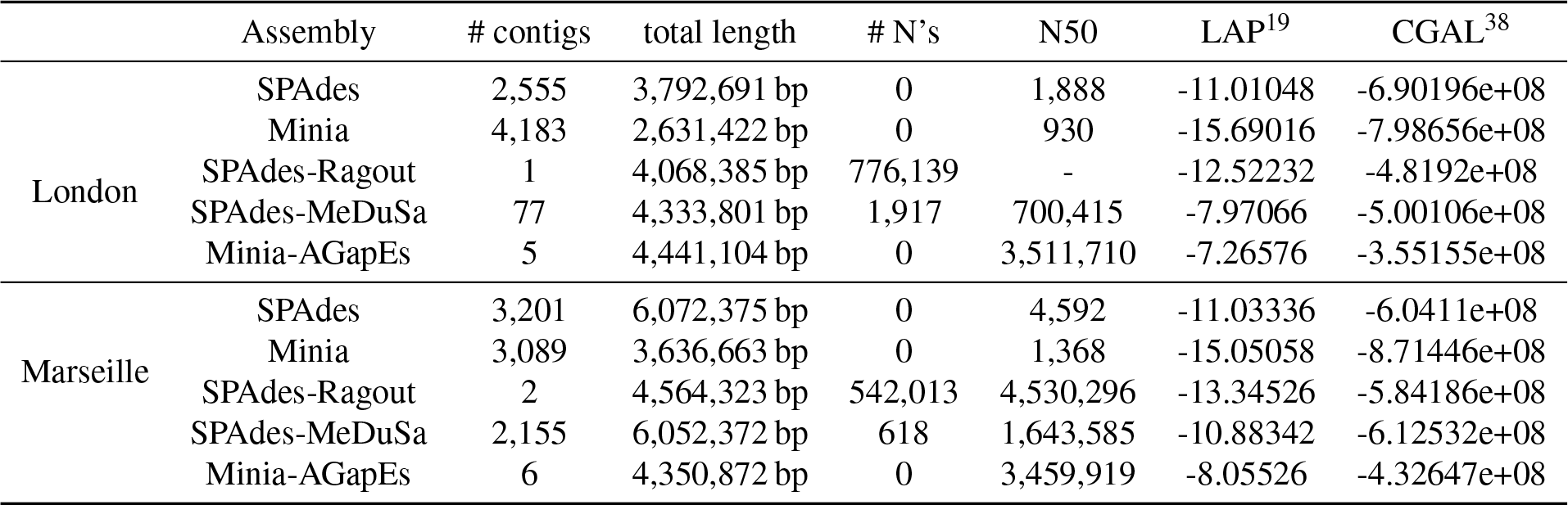
Assembly statistics for both data sets, based on contigs with a minimal length of 500*bp*. The LAP and CGAL likelihoods have been computed based on all reads mapping to any of the reference sequences. Ragout and MeDuSa depend on the quality of the initial assembly in terms of assembled sequence length, hence we omit results for the minia assembly here and refer to suppl. material Table S10.

|  | Assembly | # contigs | total length | # N's | N50 | LAP<sup>19</sup> | CGAL<sup>38</sup> |
| --- | --- | --- | --- | --- | --- | --- | --- |
| London | SPAdes | 2,555 | 3,792,691 bp | 0 | 1,888 | -11.01048 | -6.90196e+08 |
|  | Minia | 4,183 | 2,631,422 bp | 0 | 930 | -15.69016 | -7.98656e+08 |
|  | SPAdes-Ragout | 1 | 4,068,385 bp | 776,139 | - | -12.52232 | -4.8192e+08 |
|  | SPAdes-MeDuSa | 77 | 4,333,801 bp | 1,917 | 700,415 | -7.97066 | -5.00106e+08 |
|  | Minia-AGapEs | 5 | 4,441,104 bp | 0 | 3,511,710 | -7.26576 | -3.55155e+08 |
| Marseille | SPAdes | 3,201 | 6,072,375 bp | 0 | 4,592 | -11.03336 | -6.0411e+08 |
|  | Minia | 3,089 | 3,636,663 bp | 0 | 1,368 | -15.05058 | -8.71446e+08 |
|  | SPAdes-Ragout | 2 | 4,564,323 bp | 542,013 | 4,530,296 | -13.34526 | -5.84186e+08 |
|  | SPAdes-MeDuSa | 2,155 | 6,052,372 bp | 618 | 1,643,585 | -10.88342 | -6.12532e+08 |
|  | Minia-AGapEs | 6 | 4,350,872 bp | 0 | 3,459,919 | -8.05526 | -4.32647e+08 |

#### IS reconstruction

In order to validate the IS reconstruction in our assemblies, we ran the tool ISseeker^1^ that allows to annotate IS elements in draft genome assemblies by blasting flanking sequences against a reference. We tested both SPAdes and minia assemblies for the presence of 10 *Yersinia pestis* species-specific IS elements found in the ISFinder database^45^ and using all potential IS gaps as references.

While ISseeker is not able to annotate IS elements in the Minia assembly, we see in Table 2 30 annotations that are found in the SPAdes assembly. Seven of these are not annotated in the AGapEs reconstruction, and they all concern gaps that are only partially covered by reads. However a manual check of these gaps determined the presence of the respective IS element in five gaps, indicating that ISseeker was not able to correctly annotate these elements in these cases.

**Table 2.** IS annotations in London dataset by ISseeker in either draft assembly, AGapEs reconstruction or both.

|  | SPAdes | AGapEs | both | Minia | AGapEs | both |
| --- | --- | --- | --- | --- | --- | --- |
| IS gap template | 7 | 55 | 23 | 0 | 78 | 0 |

### 3.5 Discussion

In this paper, we present a method to fill the gaps between contigs assembled from aDNA reads that combines comparative scaffolding using related extant genomes and direct aDNA sequencing data, and we apply it to two ancient *Yersinia pestis* strains isolated from the remains of victims of the second plague pandemic.

The comparison of the two assemblies for the London strain illustrates that relying on a shorter initial de novo contig assembly does not impact significantly the final result. The results we obtain with the Marseille data set illustrates that if a good coverage of reads over the whole genome can be provided (as through multiple sequencing experiments for multiple samples), even a cautious initial contig assembly can be improved in such a way that most gaps are filled using unassembled aDNA reads. With both data sets, we obtain largely improved genome assemblies, with a reduced fragmentation (from thousand of contigs to a handful of CARs) and a very small fraction of the final assembly that is not supported by aDNA reads.

Applied to the same data set for the London strain, the method FPSAC^39^ was able to obtain a single scaffold based on parsimonious optimization. Comparing our resulting assembly to this single scaffold, we can identify two breakpoints between both assemblies, hence both methods do not entirely support the same scaffold structure for the London strain. These disagreements should be seen as weak points in both assemblies, as they are not reconstructed by different scaffolding objectives and would need to be confirmed more confidently by additional sequencing data.

We see a clear connection between conflicts in the set of potential adjacencies and the presence of IS elements in the corresponding gaps. Solving these conflicts based on aDNA read data provides a useful way to identify ancestral adjacencies in a conflicting component if the quality of the aDNA data is sufficient. The mapping of aDNA reads has shown to be mostly difficult at repetitive regions like Insertion Sequences, where the presence of the IS in the ancestral gap cannot be clearly detected by the aDNA sequencing data.

Interestingly, the improved assemblies of the London and Marseille strains show no explicit large genome rearrangements except for a small inversion. Even if potential genome rearrangement might not be observed due to the fragmentation of the assemblies into CARs, the synteny conservation between two strains separated by roughly 400 years of evolution is striking compared to the level of syntenic divergence with extant strains. This might be explained by the fact that both the London and Marseille strains belong to a relatively localized, although long-lasting, pandemic^6^. Also of interest is the observation that conflicting adjacencies in the Marseille data set were covered by aDNA reads, thus making it difficult to infer robust scaffolding adjacencies; this raises the question of the presence of several strains in the Marseille pandemic that might have differed by one or a few inversions.

Answering these questions with confidence would require additional targeted sequencing of a few regions of the genomes of the London and Marseille strains, or the sequencing of additional strains of the second plague pandemic, such as the *Yersinia pestis* genome sequenced from plague victims in Ellwangen^46^ which is assumed to be an ancestor of the Marseille strains.

## Authors contribution

C. C. designed the study. N. L. and C. C. designed the AGapEs method. N. L. implemented the method and analyzed data. D. D. designed and implemented the IS annotation method. C. C. and N. L. wrote the manuscript. All authors read and approved the paper.

## Acknowledements

We thank the group of Hendrik Poinar for help with the processing of the raw reads and discussions during a visit in Hamilton.

